## Supplementary Table S1 for "Mortality of native and invasive ladybirds co-infected by ectoparasitic and entomopathogenic fungi"

**Supplementary Tables**

**Table S1.** Effects of different treatments on ladybird mortality, with standard error.

| Ladybird | *He. virescens* | Treatment | Average mortality [%] | Std. Error |
|---|---|---|---|---|
| *Harmonia axyridis* | *He. virescens*-negative | Control | 5.6 | 4.14 |
| | | GHA Bb | 5.7 | 2.78 |
| | | Native Bb | 5.2 | 2.74 |
| | | Ma | 22.5 | N/A |
| | *He. virescens*-positive | Control | 53.6 | 14.21 |
| | | GHA Bb | 44.9 | 15.06 |
| | | Native Bb | 53.6 | 6.53 |
| | | Ma | 60.0 | N/A |
| *Olla v-nigrum* | *He. virescens*-negative | Control | 4.1 | 1.84 |
| | | GHA Bb | 35.3 | 20.14 |
| | | Native Bb | 57.5 | 10.45 |
| | | Ma | 60.0 | N/A |
| | *He. virescens*-positive | Control | 35.7 | 6.91 |
| | | GHA Bb | 68.4 | 9.75 |
| | | Native Bb | 91.0 | 2.85 |
| | | Ma | 97.4 | N/A |

**Table S2.** Overview of number of ladybirds used per treatment. Presented are numbers for the different replicates (separated by "+") within each assay.

| **Ladybird** | ***He. virescens*** | **Treatment** | **Assay #1** | **Assay #2** | **Assay #3** |
|---|---|---|---|---|---|
| *Harmonia axyridis* | No | Control | 6+6+6 | 10+9+10 | 10+10+10+10 |
| | | GHA Bb | 6+6+6 | 10+10+10 | 10+10+10+10 |
| | | Native Bb | 6+6+6 | 10+10+10 | 10+10+9+10 |
| | | Ma | N/A | N/A | 10+10+10+10 |
| | Yes | Control | 10+10+10 | 8+8+8 | 10+9+10+10 |
| | | GHA Bb | 10+10+10 | 8+8+7 | 10+10+10+10 |
| | | Native Bb | 10+10+10 | 8+8+8 | 10+10+10+10 |
| | | Ma | N/A | N/A | 10+10+10+10 |
| *Olla v-nigrum* | No | Control | 10+10+10 | 10+10+10 | 10+10+10+10 |
| | | GHA Bb | 10+10+10 | 10+10+10 | 10+10+10+10 |
| | | Native Bb | 10+10+10 | 10+10+10 | 10+9+10+10 |
| | | Ma | N/A | N/A | 10+10+10+10 |
| | Yes | Control | 10+10+10 | 9+9+9 | 10+10+10+10 |
| | | GHA Bb | 10+10+9 | 9+9+9 | 10+10+10+10 |
| | | Native Bb | 10+10+10 | 9+9+9 | 10+10+10+10 |
| | | Ma | N/A | N/A | 10+8+10+10 |
